# North America Insect Invades the World: the Case of the Flood Mosquito, *Aedes vexans* (Diptera: Culicidae), from Mitochondrial DNA

**DOI:** 10.1101/2021.04.15.439950

**Authors:** José Heriberto Vargas-Espinosa, Oscar Alexander Aguirre-Obando

## Abstract

The flood mosquito, *Aedes vexans* (Diptera: Culicidae), native of Canada, and currently present in all continents, has a vector competence for 30 arboviruses, being responsible for transmitting diseases, like West Nile fever, Rift Valley fever, Saint Louis Encephalitis and Eastern Equine Encephalitis. Hence, knowing the structure and gene flow of *A. vexans* is important to develop adequate vector control strategies for this species. For this, from partial sequences of the mitochondrial COI gene available in Bold and GenBank, it was possible to determine the Haplotypic (*Hd*) and nucleotide (π) gene diversity, genetic structuring and gene flow at global, continental, and country levels. In total, 1184 sequences were recovered, distributed between America (88.60%), Europe (7.35%), Asia (3.89%), and Africa (0.17%). From these, 395 haplotypes (H) were detected without presence of pseudogenes (NUMTs), with H1 being the most frequent (24.58%) and between H12 - H395 the least frequent varying between 0.93% (H12) and 0.08% (H395). Phylogenetically, the haplotypes were grouped into six clades. Clade I grouped haplotypes from countries in America and Europe, while clades II and III presented haplotypes exclusively from Asia and Europe; clade IV grouped only one haplotype from Africa and the last ultimo clade V grouped haplotypes from America and Africa. The global *Hd* and π was 0.92 and 0.01, respectively. In addition, evidence was obtained of genetic structuring among continents (7.07%), countries (1.62%), and within countries (91.30%; F_ST_ = 0.08, p < 0.05) and no isolation by distance was detected (r = 0.003, p > 0.05). These results suggest that the mosquito populations that invaded other continents originate directly from the American continent, where possibly transcontinental commercial routes favored their long-distance dispersion.

## Introduction

The flood mosquito, *Aedes vexans* (Meigen, 1830), is present in subtropical regions of all continents, except for the Antarctic [1,2,3]. In nature this species manages to travel up to 17 km [4]; however, its global invasion was favored through passive transport mediated by human activities [5]. Like other mosquitos of medical and veterinary importance [6], *A. vexans* lays its eggs in moist sites with flood probability to guarantee their offspring [7] and these can resist drying out and survive up to three years when kept moist [8]. In general, *A. vexans* females and males feed on nectar, but to mature their ovaries and reproduce, the females feed principally on mammals, like deer, horses, cattle and pigs [9,10,11]. Normally, this species is found in rural zones, but when it inhabits suburban and urban zones, it prefers humans as principal feeding source [12,13].

Morphological and molecular evidence suggest the existence of three subspecies of *A. vexans* throughout the world: *Aedes vexans vexans* Meigen in eastern Asia and Oceania, *Aedes vexans arabiensis* Patton in Africa and Europe, and *Aedes vexans nipponii* Theobald in southeast Asia [1,14,15,16,17]. The flood mosquito has vector competence for 30 arbovirus (Elizondo-Quiroga *et al*. 2018) and is involved in the transmission of important diseases, like the West Nile fever, Rift Valley fever, Saint Louis encephalitis and Eastern Equine encephalitis, as well as filarial nematodes [18,19].

Given the epidemiological and sanitary importance of *A. vexans*, understanding the structuring patterns and gene flow of this species’ populations is important to develop more-adequate vector control programs [20], as well as understand the transmission of vectors among the human population, given its influence on pathogen transfer and dissemination of characteristics, like vector competence and resistance to insecticides [21,22]. For example, studies on genetic structure in populations of *Aedes aegypti*, dengue, chikungunya, and Zika vector, in two locations of Queensland, Australia, indicated that said locations were partially isolated genetically, with these two sites being adequate for the release of mosquitos infected with *Wolbachia pipientis* because it was important to restrict the strain released during the initial implementation phases [23,24]. Thereafter, mosquitos infected with *Wolbachia pipientis* were released with successful establishment in both locations, thus, suppressing dengue transmission [25].

Molecular markers are widely used to understand the biology and population dynamics of vector species of diseases [26]. Among the molecular markers used in population genetics studies in *A. vexans*, the mitochondrial DNA (mtDNA) sequences are broadly used due to properties, like their abundance in the organism, size and small genomic structure, rapid rate of evolution, and exclusive maternal inheritance with low genetic recombination [27]. However, one of the disadvantages of using mtDNA in population genetics and phylogenetic studies is NUMTs presence, result of the translocation of mitochondrial sequences of the mitochondrial genome for the nuclear genome [28]. Furthermore, this type of information is freely available in the GenBank and Boldsystem databases. Particularly for *A. vexans*, until now, nobody has analyzed the genetic information available within a global context. This work sought to know the global panorama of genetic diversity and gene flow of *A. vexans*, using mtDNA sequences available in GenBank and Boldsystem.

## Materials and Methods

A prior search in GenBank permitted detecting that the mtDNA Cytochrome oxidase subunit I (COI) gene was the most representative, and given that Boldsystem is this marker’s depository, it was also determined to work with it. For this, partial nucleotide sequences were obtained from the mitochondrial (mtDNA) COI gene deposited in GenBank and Boldsystem for *A. vexans*. The search criteria in GenBank used the words *Aedes vexans* AND COI, while for Boldsystem, it only used *Aedes vexans*. The sequences obtained were analyzed by using the BLAST tool in the NCBI website (http://blast.ncbi.nlm.nih.gov/Blast.cgi) to confirm identity with *A. vexans* and only sequences with identity percentage between 98% and 100% were considered in our analysis. Thereafter, the sequences selected were classified according to continent and country, so that analyses listed ahead were performed for each of them. In addition, for each sequence, geographic data were extracted to later geo-reference it on a map; sequences without geographic information were eliminated from the analysis. These data were filtered and organized through the RStudio platform by using Bold packages version 0.9 [29] and Ape version 5.3 [30].

Then, the sequences were aligned by using the MAFFT software version 7 to later detect haplotypes (H) [31]. Each haplotype was numbered based on its frequency, thus, the most frequent was H1, the following H2, and so forth. To detect potential NUMTs among the H detected, additional stop codons were searched for in the alignment [28]. In case of detecting any NUMT, it was reported and removed from the analysis. With H without NUMTs, the haplotype network was constructed through the Pegas package version 0.11 [32] in the RStudio platform [33].

Diversity and neutrality tests were estimated by using the DnaSP program version 6.0 [34]. The analysis of molecular variance (AMOVA) was conducted by using the Arlequin program version 3.5 [35], which evaluated the variation between continents and countries. Population genetic structuring was tested by using the fixation index (F_ST_) proposed by Wright [36], and the gene flow (Nm) was calculated through the Arlequin program version 3.5 [35]. In cases in which statistically significant differences were obtained, the Bonferroni correction was used to verify the existence of significant differences.

To test isolation by distance, Mantel’s test was carried out to estimate the correlation between genetic distance (FST) and geographic distance (Km) using the Vegan package version 2.5 [37] on the RStudio platform [33]. The geographic distances were obtained by calculating with the Geographic Distance Matrix Generator program version 1.2.3. To estimate the genetic affinity of *A. vexans* populations, a phylogenetic reconstruction was performed using maximum likelihood (ML) and Bayesian inference (BI) from the haplotypes found. For this, we initially searched for the nucleotide substitution model that best fits our data in the program jModelTest version 2.1.1, which selected the model with the lowest value from the Akaike information criterion, AIC, [38]. Then, the model selected was used for the phylogenetic reconstruction under the ML and BI approach. The ML analysis was conducted with the RAxML software [39], under the following parameters: ML+ thorough Bootstrap and 1,000 boot replicas. In turn, the BI analysis was conducted in the Mr.Bayes program version 3.2.7, under the following parameters: number of generations = 2,000,000, with σ < 0.01 of the frequencies to indicate robustness of the phylogenetic hypothesis [40]. The visualization and editing of the phylogenetic trees obtained was carried out in Mr.Ent version 2.5 [41].

## Results

Between April and May 2020, the study recovered 2,420 sequences from the Boldsystem (82.64%) and GenBank (17.35%) databases, distributed among America (94.50%), Europe (3.68%), Asia (1.23%), and Africa (0.58%) with median length of 467 bp, varying between 114 and 879 bp. Nevertheless, once the alignment was made, 1,184 sequences were selected from these with a length of 340 bp each, all distributed among the continents mentioned. Among the sequences selected, the American continent had the highest representation (88.60%), followed by Europe (7.35%), Asia (3.89%), and Africa (0.17%). The American continent was represented with sequences from Canada (64.10%) and the United States (24.49%). Europe was represented by Sweden (2.87%), Belgium (1.35%), Spain (1.27%), Netherlands (0.84%), Austria (1.35%), Germany (0.17%), Rumania (0.17%), Germany (0.17%), Kosovo (0.17%), Hungary (0.17 %), and Turkey (0.17%). Asia was represented by Japan (1.10%), China (1.10%), Iran (0.59%), Russia (0.42%), Singapore (0.25%), South Korea (0.25%), and India (0.17%). Finally, Africa was represented only by South Africa (0.17%).

Table 1 shows the global distribution of the haplotypes observed. In total, 395 H were observed, with H1 being the most frequent (24.58%), followed by H2 (7.77%), H3 (4.39%), H4 (3.38%), H5 (2.28%), H6 - H11 ranging between 1.86% (H6) and 1.10% (H11), and H112-H395 varying between 0.93% (H112) and 0.08% (H395). Although H1 was the most frequent, it was only observed in Canada and the USA. Nevertheless, the haplotype with the greatest distribution was H7, which was present in eight countries, that is, Austria (11.76%), Belgium (17.64%), Hungary (11.76%), Kosovo (5.88%), Netherlands (17.64%), Russia (5.88%), Spain (11.76%), and Sweden (17.64%). None of the H had presence of NUMTs (Supplementary material). Figure 1 displays the locations where the genetic material analyzed was extracted (Fig. 1A), as well as the phylogenetic relationships between populations, as arrows observed in the haplotype network (Fig. 1B). From this, it may be stated that native populations of *A. vexans* from America invaded some countries in Africa and Europe, and from there established themselves in other countries in these continents and in Asia.

| Haplotype | Continent |  |  |  | Access code | Reference |
| --- | --- | --- | --- | --- | --- | --- |
|  | America | Africa | Asia | Europe |  |  |
| H1 | 291 | 0 | 0 | 0 | ACMC131-04 | [42] |
| H2 | 92 | 0 | 0 | 0 | ACMC277-04 | [42] |
| H3 | 52 | 0 | 0 | 0 | CNEIF006-12 | [43] |
| H4 | 40 | 0 | 0 | 0 | ACMC276-04 | [42] |
| H5 | 27 | 0 | 0 | 0 | CNGSD2112-15 | [43] |
| H6 | 0 | 0 | 0 | 22 | CULBE143-14 | [44] |
| H7 | 0 | 0 | 0 | 17 | GBMIN56980-17 | [44] |
| H8 | 15 | 0 | 0 | 0 | ASDIP113-15 | http://www.boldsystems.org/ |
| H9 | 13 | 0 | 0 | 0 | ACMC289-04 | [42] |
| H10 | 12 | 0 | 0 | 0 | BBDCP596-10 | http://www.boldsystems.org/ |
| H11 | 13 | 0 | 0 | 0 | CNPAD261-13 | [43] |
| H12 | 11 | 0 | 0 | 0 | CNEIF841-12 | [43] |
| H13 | 10 | 0 | 0 | 0 | CNEIF861-12 | [43] |
| H14 | 9 | 0 | 0 | 0 | ASDMT244-11 | [43] |
| H15 | 9 | 0 | 0 | 0 | ACMC290-04 | [42] |
| H16 | 7 | 0 | 0 | 0 | CNGSD6857-15 | [43] |
| H17 | 7 | 0 | 0 | 0 | CNGSD860-15 | [43] |
| H18 | 7 | 0 | 0 | 0 | CNJAE1246-12 | [43] |
| H19 | 7 | 0 | 0 | 0 | CNPAF983-13 | [43] |
| H20 | 7 | 0 | 0 | 0 | SSJAE8713-13 | [43] |
| H21 | 6 | 0 | 0 | 0 | CNJAE1248-12 | [43] |
| H22 | 0 | 0 | 8 | 0 | GBDP47430-19 | [45] |
| H23 | 5 | 0 | 0 | 0 | CNEIF812-12 | [43] |
| H24 | 5 | 0 | 0 | 0 | MBIOI074-13 | [43] |
| H25 | 5 | 0 | 0 | 0 | RRSSA3576-15 | [49] |
| H26 | 5 | 0 | 0 | 0 | SSEIC6245-13 | [43] |
| H27 | 5 | 0 | 0 | 0 | SSJAE8835-13 | [43] |
| H28 | 0 | 0 | 0 | 4 | CULBE132-14 | [44] |
| H29 | 4 | 0 | 0 | 0 | ACMC104-04 | [42] |
| H30 | 4 | 0 | 0 | 0 | CNEIG153-12 | [43] |
| H31 | 4 | 0 | 0 | 0 | CNGSD1517-15 | [43] |
| H32 | 4 | 0 | 0 | 0 | CNPAD244-13 | [43] |
| H33 | 4 | 0 | 0 | 0 | PHJUL2502-11 | [43] |
| H34 | 4 | 0 | 0 | 0 | SSEIC5403-13 | [43] |
| H35 | 4 | 0 | 0 | 0 | SSEIC5955-13 | [43] |
| H36 | 4 | 0 | 0 | 0 | SSEIC6896-13 | [43] |
| H37 | 4 | 0 | 0 | 0 | SSJAE8703-13 | [43] |
| H38 | 4 | 0 | 0 | 0 | SSJAE9027-13 | [43] |
| H39 | 0 | 0 | 0 | 4 | GBDP48120-19 | [48] |
| H40 | 3 | 0 | 0 | 0 | CNEIF014-12 | [43] |
| H41 | 3 | 0 | 0 | 0 | CNEIF817-12 | [43] |
| H42 | 3 | 0 | 0 | 0 | CNEIF830-12 | [43] |
| H43 | 3 | 0 | 0 | 0 | CNEIG182-12 | [43] |
| H44 | 3 | 0 | 0 | 0 | CNGSI1668-15 | [43] |
| H45 | 3 | 0 | 0 | 0 | CNRME208-12 | [43] |
| H46 | 3 | 0 | 0 | 0 | PHAUG1312-11 | [43] |
| H47 | 3 | 0 | 0 | 0 | SMTPJ4980-14 | [43] |
| H48 | 3 | 0 | 0 | 0 | SSEIA4548-13 | [43] |
| H49 | 3 | 0 | 0 | 0 | SSEIA5279-13 | [43] |
| H50 | 3 | 0 | 0 | 0 | SSEIA5341-13 | [43] |
| H51 | 3 | 0 | 0 | 0 | SSJAF9177-13 | [43] |
| H52 | 3 | 0 | 0 | 0 | SSPAC12518-13 | [43] |
| H53 | 3 | 0 | 0 | 0 | SSPPA9368-15 | [43] |
| H54 | 0 | 0 | 3 | 0 | MK962499 | [50] |
| H55 | 0 | 0 | 3 | 0 | AB738099 | [45] |
| H56 | 0 | 0 | 0 | 3 | GBMIN56959-17 | [48] |
| H57 | 3 | 0 | 0 | 0 | IUP1017-14 | http://www.boldsystems.org/ |
| H58 | 0 | 0 | 0 | 2 | CULBE140-14 | [44] |
| H59 | 2 | 0 | 0 | 0 | ACMC033-04 | [42] |
| H60 | 2 | 0 | 0 | 0 | BBDCM210-10 | [43] |
| H61 | 2 | 0 | 0 | 0 | CNEIF002-12 | [43] |
| H62 | 2 | 0 | 0 | 0 | CNEIF798-12 | [43] |
| H63 | 2 | 0 | 0 | 0 | CNGSE2411-15 | [43] |
| H64 | 2 | 0 | 0 | 0 | CNGSI1097-15 | [43] |
| H65 | 2 | 0 | 0 | 0 | CNPKO3332-14 | [43] |
| H66 | 2 | 0 | 0 | 0 | CNRME207-12 | [43] |
| H67 | 2 | 0 | 0 | 0 | CNRME2449-12 | [43] |
| H68 | 2 | 0 | 0 | 0 | CNRME4093-12 | [43] |
| H69 | 2 | 0 | 0 | 0 | CNRMG626-12 | [43] |
| H70 | 2 | 0 | 0 | 0 | CNTIH462-15 | [43] |
| H71 | 2 | 0 | 0 | 0 | GMOKH968-15 | [43] |
| H72 | 2 | 0 | 0 | 0 | SSEIA3424-13 | [43] |
| H73 | 2 | 0 | 0 | 0 | SSEIA3430-13 | [43] |
| H74 | 2 | 0 | 0 | 0 | SSEIA5215-13 | [43] |
| H75 | 2 | 0 | 0 | 0 | SSEIA5266-13 | [43] |
| H76 | 2 | 0 | 0 | 0 | SSJAE1181-13 | [43] |
| H77 | 2 | 0 | 0 | 0 | SSJAE8706-13 | [43] |
| H78 | 2 | 0 | 0 | 0 | SSJAF9208-13 | [43] |
| H79 | 2 | 0 | 0 | 0 | SSPAB8912-13 | [43] |
| H80 | 2 | 0 | 0 | 0 | SSPAC12020-13 | [43] |
| H81 | 2 | 0 | 0 | 0 | SSPPA5271-15 | [43] |
| H82 | 2 | 0 | 0 | 0 | SSPPA5483-15 | [43] |
| H83 | 2 | 0 | 0 | 0 | SSPPA9400-15 | [43] |
| H84 | 0 | 0 | 2 | 0 | GBDP47423-19 | [45] |
| H85 | 0 | 0 | 2 | 0 | AB738168- | [45] |
| H86 | 0 | 0 | 0 | 2 | GBMIN56950-17 | [46] |
| H87 | 0 | 0 | 1 | 1 | LRMQS005-17 | [47] |
| H88 | 0 | 0 | 1 | 1 | GBDP48149-19 | [48] |
| H89 | 0 | 0 | 1 | 1 | GBDP48185-19 | [48] |
| H90 | 0 | 0 | 1 | 1 | GBDP48490-19 | [48] |
| H91 | 0 | 0 | 0 | 2 | GBMIN56961-17 | [48] |
| H92 | 2 | 0 | 0 | 0 | MOSN025-17 | http://www.boldsystems.org/ |
| H93 | 2 | 0 | 0 | 0 | MOSN050-17 | http://www.boldsystems.org/ |
| H94 | 2 | 0 | 0 | 0 | MOSN1411-18 | http://www.boldsystems.org/ |
| H95 | 1 | 0 | 0 | 0 | MOSN1957-18 | http://www.boldsystems.org/ |
| H96 | 1 | 0 | 0 | 0 | MOSN2259-19 | http://www.boldsystems.org/ |
| H97 | 1 | 0 | 0 | 0 | MOSN2970-19 | http://www.boldsystems.org/ |
| H98 | 0 | 0 | 0 | 1 | CULBE141-14 | [44] |
| H99 | 0 | 0 | 0 | 1 | CULBE145-14 | [44] |
| H100 | 0 | 0 | 0 | 1 | GMOKH2148-15 | [43] |
| H101 | 0 | 0 | 0 | 1 | CULBE136-14 | [44] |
| H102 | 0 | 0 | 0 | 1 | CULBE138-14 | [44] |
| H103 | 0 | 0 | 0 | 1 | CULBE139-14 | [44] |
| H104 | 1 | 0 | 0 | 0 | ACMC053-04 | [42] |
| H105 | 1 | 0 | 0 | 0 | ACMC288-04 | [42] |
| H106 | 1 | 0 | 0 | 0 | ASDMT1059-11 | [43] |
| H107 | 1 | 0 | 0 | 0 | BBDCN133-10 | [43] |
| H108 | 1 | 0 | 0 | 0 | BBDCN233-10 | [43] |
| H109 | 1 | 0 | 0 | 0 | BBDCP593-10 | [43] |
| H110 | 1 | 0 | 0 | 0 | CNEIF015-12 | [43] |
| H111 | 1 | 0 | 0 | 0 | CNEIF023-12 | [43] |
| H112 | 1 | 0 | 0 | 0 | CNEIF3339-12 | [43] |
| H113 | 1 | 0 | 0 | 0 | CNEIF793-12 | [43] |
| H114 | 1 | 0 | 0 | 0 | CNEIF813-12 | [43] |
| H115 | 1 | 0 | 0 | 0 | CNEIF849-12 | [43] |
| H116 | 1 | 0 | 0 | 0 | CNEIG1456-12 | [43] |
| H117 | 1 | 0 | 0 | 0 | CNEIG152-12 | [43] |
| H118 | 1 | 0 | 0 | 0 | CNFNH1033-14 | [43] |
| H119 | 1 | 0 | 0 | 0 | CNGRD1138-12 | [43] |
| H120 | 1 | 0 | 0 | 0 | CNGSD1018-15 | [43] |
| H121 | 1 | 0 | 0 | 0 | CNGSD132-15 | [43] |
| H122 | 1 | 0 | 0 | 0 | CNGSD1611-15 | [43] |
| H123 | 1 | 0 | 0 | 0 | CNGSD263-15 | [43] |
| H124 | 1 | 0 | 0 | 0 | CNGSE1406-15 | [43] |
| H125 | 1 | 0 | 0 | 0 | CNGSE2070-15 | [43] |
| H126 | 1 | 0 | 0 | 0 | CNGSE2173-15 | [43] |
| H127 | 1 | 0 | 0 | 0 | CNGSG1207-15 | [43] |
| H128 | 1 | 0 | 0 | 0 | CNGSG122-15 | [43] |
| H129 | 1 | 0 | 0 | 0 | CNGSI1101-15 | [43] |
| H130 | 1 | 0 | 0 | 0 | CNGSI1218-15 | [43] |
| H131 | 1 | 0 | 0 | 0 | CNGSI1988-15 | [43] |
| H132 | 1 | 0 | 0 | 0 | CNGSI246-15 | [43] |
| H133 | 1 | 0 | 0 | 0 | CNGSI326-15 | [43] |
| H134 | 1 | 0 | 0 | 0 | CNGSI582-15 | [43] |
| H135 | 1 | 0 | 0 | 0 | CNJAE1277-12 | [43] |
| H136 | 1 | 0 | 0 | 0 | CNJAE965-12 | [43] |
| H137 | 1 | 0 | 0 | 0 | CNJAF2111-12 | [43] |
| H138 | 1 | 0 | 0 | 0 | CNJAF2114-12 | [43] |
| H139 | 1 | 0 | 0 | 0 | CNJAF2142-12 | [43] |
| H140 | 1 | 0 | 0 | 0 | CNPAD242-13 | [43] |
| H141 | 1 | 0 | 0 | 0 | CNPAD260-13 | [43] |
| H142 | 1 | 0 | 0 | 0 | CNPPC1400-12 | [43] |
| H143 | 1 | 0 | 0 | 0 | CNPPC1722-12 | [43] |
| H144 | 1 | 0 | 0 | 0 | CNRME198-12 | [43] |
| H145 | 1 | 0 | 0 | 0 | CNRME2450-12 | [43] |
| H146 | 1 | 0 | 0 | 0 | CNRMG169-12 | [43] |
| H147 | 1 | 0 | 0 | 0 | CNROH071-13 | [43] |
| H148 | 1 | 0 | 0 | 0 | CNTIH336-15 | [43] |
| H149 | 1 | 0 | 0 | 0 | CNWBC188-13 | [43] |
| H150 | 1 | 0 | 0 | 0 | CNWBH228-13 | [43] |
| H151 | 1 | 0 | 0 | 0 | CNWBH672-13 | [43] |
| H152 | 1 | 0 | 0 | 0 | GMOKH2148-15 | [43] |
| H153 | 1 | 0 | 0 | 0 | JSJUN1501-11 | [43] |
| H154 | 1 | 0 | 0 | 0 | MSQ002-04 | http://www.boldsystems.org/ |
| H155 | 1 | 0 | 0 | 0 | MSQ003-04 | http://www.boldsystems.org/ |
| H156 | 1 | 0 | 0 | 0 | MSQ015-04 | http://www.boldsystems.org/ |
| H157 | 1 | 0 | 0 | 0 | MSQ022-04 | http://www.boldsystems.org/ |
| H158 | 1 | 0 | 0 | 0 | NCCA889-11 | [43] |
| H159 | 1 | 0 | 0 | 0 | NCCA895-11 | [43] |
| H160 | 1 | 0 | 0 | 0 | PHAUG1277-11 | [43] |
| H161 | 1 | 0 | 0 | 0 | PHAUG1318-11 | [43] |
| H162 | 1 | 0 | 0 | 0 | PHSEP1211-11 | [43] |
| H163 | 1 | 0 | 0 | 0 | PREXP286-14 | http://www.boldsystems.org/ |
| H164 | 1 | 0 | 0 | 0 | RRINV213-15 | [49] |
| H165 | 1 | 0 | 0 | 0 | RRINV3712-15 | [49] |
| H166 | 1 | 0 | 0 | 0 | RRINV3721-15 | [49] |
| H167 | 1 | 0 | 0 | 0 | RRINV3723-15 | [49] |
| H168 | 1 | 0 | 0 | 0 | RRINV3798-15 | [49] |
| H169 | 1 | 0 | 0 | 0 | RRMFI4275-15 | [49] |
| H170 | 1 | 0 | 0 | 0 | RRSSA425-15 | [49] |
| H171 | 1 | 0 | 0 | 0 | SMTPI2660-14 | [43] |
| H172 | 1 | 0 | 0 | 0 | SMTPI3303-14 | [43] |
| H173 | 1 | 0 | 0 | 0 | SSBAE5601-13 | [43] |
| H174 | 1 | 0 | 0 | 0 | SSBAE5713-13 | [43] |
| H175 | 1 | 0 | 0 | 0 | SSEIA3425-13 | [43] |
| H176 | 1 | 0 | 0 | 0 | SSEIA4525-13 | [43] |
| H177 | 1 | 0 | 0 | 0 | SSEIA5135-13 | [43] |
| H178 | 1 | 0 | 0 | 0 | SSEIA5184-13 | [43] |
| H179 | 1 | 0 | 0 | 0 | SSEIA5193-13 | [43] |
| H180 | 1 | 0 | 0 | 0 | SSEIA5225-13 | [43] |
| H181 | 1 | 0 | 0 | 0 | SSEIA5296-13 | [43] |
| H182 | 1 | 0 | 0 | 0 | SSEIA5301-13 | [43] |
| H183 | 1 | 0 | 0 | 0 | SSEIA5335-13 | [43] |
| H184 | 1 | 0 | 0 | 0 | SSEIC5215-13 | [43] |
| H185 | 1 | 0 | 0 | 0 | SSEIC5397-13 | [43] |
| H186 | 1 | 0 | 0 | 0 | SSEIC5788-13 | [43] |
| H187 | 1 | 0 | 0 | 0 | SSEIC5981-13 | [43] |
| H188 | 1 | 0 | 0 | 0 | SSEIC6590-13 | [43] |
| H189 | 1 | 0 | 0 | 0 | SSEIC6788-13 | [43] |
| H190 | 1 | 0 | 0 | 0 | SSEIC6792-13 | [43] |
| H191 | 1 | 0 | 0 | 0 | SSEIC6946-13 | [43] |
| H192 | 1 | 0 | 0 | 0 | SSEID823-13 | [43] |
| H193 | 1 | 0 | 0 | 0 | SSJAE10960-13 | [43] |
| H194 | 1 | 0 | 0 | 0 | SSJAE11005-13 | [43] |
| H195 | 1 | 0 | 0 | 0 | SSJAE5456-13 | [43] |
| H196 | 1 | 0 | 0 | 0 | SSJAE6982-13 | [43] |
| H197 | 1 | 0 | 0 | 0 | SSJAE7467-13 | [43] |
| H198 | 1 | 0 | 0 | 0 | SSJAE8697-13 | [43] |
| H199 | 1 | 0 | 0 | 0 | SSJAE8710-13 | [43] |
| H200 | 1 | 0 | 0 | 0 | SSJAE8721-13 | [43] |
| H201 | 1 | 0 | 0 | 0 | SSJAE8767-13 | [43] |
| H202 | 1 | 0 | 0 | 0 | SSJAE8770-13 | [43] |
| H203 | 1 | 0 | 0 | 0 | SSJAE8773-13 | [43] |
| H204 | 1 | 0 | 0 | 0 | SSJAE8790-13 | [43] |
| H205 | 1 | 0 | 0 | 0 | SSJAE8802-13 | [43] |
| H206 | 1 | 0 | 0 | 0 | SSJAE8804-13 | [43] |
| H207 | 1 | 0 | 0 | 0 | SSJAE8810-13 | [43] |
| H208 | 1 | 0 | 0 | 0 | SSJAE8842-13 | [43] |
| H209 | 1 | 0 | 0 | 0 | SSJAE8865-13 | [43] |
| H210 | 1 | 0 | 0 | 0 | SSJAE8905-13 | [43] |
| H211 | 1 | 0 | 0 | 0 | SSJAE8909-13 | [43] |
| H212 | 1 | 0 | 0 | 0 | SSJAE8962-13 | [43] |
| H213 | 1 | 0 | 0 | 0 | SSJAE8972-13 | [43] |
| H214 | 1 | 0 | 0 | 0 | SSJAE8997-13 | [43] |
| H215 | 1 | 0 | 0 | 0 | SSJAE9009-13 | [43] |
| H216 | 1 | 0 | 0 | 0 | SSJAE9068-13 | [43] |
| H217 | 1 | 0 | 0 | 0 | SSJAF10000-13 | [43] |
| H218 | 1 | 0 | 0 | 0 | SSJAF10014-13 | [43] |
| H219 | 1 | 0 | 0 | 0 | SSJAF10300-13 | [43] |
| H220 | 1 | 0 | 0 | 0 | SSJAF10307-13 | [43] |
| H221 | 1 | 0 | 0 | 0 | SSJAF10358-13 | [43] |
| H222 | 1 | 0 | 0 | 0 | SSJAF10367-13 | [43] |
| H223 | 1 | 0 | 0 | 0 | SSJAF259-13 | [43] |
| H224 | 1 | 0 | 0 | 0 | SSJAF5420-13 | [43] |
| H225 | 1 | 0 | 0 | 0 | SSJAF9159-13 | [43] |
| H226 | 1 | 0 | 0 | 0 | SSJAF9168-13 | [43] |
| H227 | 1 | 0 | 0 | 0 | SSJAF9715-13 | [43] |
| H228 | 1 | 0 | 0 | 0 | SSJAF9963-13 | [43] |
| H229 | 1 | 0 | 0 | 0 | SSPAA9388-13 | [43] |
| H230 | 1 | 0 | 0 | 0 | SSPAA9453-13 | [43] |
| H231 | 1 | 0 | 0 | 0 | SSPAA9670-13 | [43] |
| H232 | 1 | 0 | 0 | 0 | SSPAA9697-13 | [43] |
| H233 | 1 | 0 | 0 | 0 | SSPAA9698-13 | [43] |
| H234 | 1 | 0 | 0 | 0 | SSPAC11523-13 | [43] |
| H235 | 1 | 0 | 0 | 0 | SSPAC11881-13 | [43] |
| H236 | 1 | 0 | 0 | 0 | SSPAC12030-13 | [43] |
| H237 | 1 | 0 | 0 | 0 | SSPAC12040-13 | [43] |
| H238 | 1 | 0 | 0 | 0 | SSPAC12053-13 | [43] |
| H239 | 1 | 0 | 0 | 0 | SSPAC12068-13 | [43] |
| H240 | 1 | 0 | 0 | 0 | SSPAC12075-13 | [43] |
| H241 | 1 | 0 | 0 | 0 | SSPAC12104-13 | [43] |
| H242 | 1 | 0 | 0 | 0 | SSPAC12117-13 | [43] |
| H243 | 1 | 0 | 0 | 0 | SSPAC12132-13 | [43] |
| H244 | 1 | 0 | 0 | 0 | SSPAC12142-13 | [43] |
| H245 | 1 | 0 | 0 | 0 | SSPAC12161-13 | [43] |
| H246 | 1 | 0 | 0 | 0 | SSPAC12173-13 | [43] |
| H247 | 1 | 0 | 0 | 0 | SSPAC12503-13 | [43] |
| H248 | 1 | 0 | 0 | 0 | SSPAC1637-13 | [43] |
| H249 | 1 | 0 | 0 | 0 | SSPPA3871-15 | [43] |
| H250 | 1 | 0 | 0 | 0 | SSPPA4038-15 | [43] |
| H251 | 1 | 0 | 0 | 0 | SSPPA4045-15 | [43] |
| H252 | 1 | 0 | 0 | 0 | SSPPA4260-15 | [43] |
| H253 | 1 | 0 | 0 | 0 | SSPPA4386-15 | [43] |
| H254 | 1 | 0 | 0 | 0 | SSPPA5066-15 | [43] |
| H255 | 1 | 0 | 0 | 0 | SSPPA5187-15 | [43] |
| H256 | 1 | 0 | 0 | 0 | SSPPA5299-15 | [43] |
| H257 | 1 | 0 | 0 | 0 | SSPPA5324-15 | [43] |
| H258 | 1 | 0 | 0 | 0 | SSPPA5593-15 | [43] |
| H259 | 1 | 0 | 0 | 0 | SSPPA5622-15 | [43] |
| H260 | 1 | 0 | 0 | 0 | SSPPA8632-15 | [43] |
| H261 | 1 | 0 | 0 | 0 | SSPPA8661-15 | [43] |
| H262 | 1 | 0 | 0 | 0 | SSPPA8754-15 | [43] |
| H263 | 1 | 0 | 0 | 0 | SSPPA8797-15 | [43] |
| H264 | 1 | 0 | 0 | 0 | SSPPA8894-15 | [43] |
| H265 | 1 | 0 | 0 | 0 | SSPPA8942-15 | [43] |
| H266 | 1 | 0 | 0 | 0 | SSPPA8947-15 | [43] |
| H267 | 1 | 0 | 0 | 0 | SSPPA9373-15 | [43] |
| H268 | 1 | 0 | 0 | 0 | SSPPA9424-15 | [43] |
| H269 | 1 | 0 | 0 | 0 | SSPPA9526-15 | [43] |
| H270 | 1 | 0 | 0 | 0 | TTDBW601-09 | [43] |
| H271 | 1 | 0 | 0 | 0 | TTMDJ190-10 | [43] |
| H272 | 0 | 0 | 1 | 0 | GBDP47418-19 | [45] |
| H273 | 0 | 0 | 1 | 0 | GBDP47419-19 | [45] |
| H274 | 0 | 0 | 1 | 0 | GBDP47420-19 | [45] |
| H275 | 0 | 0 | 1 | 0 | GBDP47421-19 | [45] |
| H276 | 0 | 0 | 1 | 0 | GBDP47422-19 | [45] |
| H277 | 0 | 0 | 1 | 0 | GBDP47424-19 | [45] |
| H278 | 0 | 0 | 1 | 0 | GBDP47425-19 | [45] |
| H279 | 0 | 0 | 1 | 0 | GBDP47427-19 | [45] |
| H280 | 0 | 0 | 1 | 0 | GBDP47428-19 | [45] |
| H281 | 0 | 0 | 1 | 0 | GBDP47429-19 | [45] |
| H282 | 0 | 0 | 0 | 1 | GBMIN56977-17 | [45] |
| H283 | 0 | 0 | 0 | 1 | GBMIN56977-18 | [49] |
| H284 | 0 | 0 | 1 | 0 | MK962500 | [50] |
| H285 | 0 | 0 | 1 | 0 | AB738152 | [45] |
| H286 | 0 | 0 | 1 | 0 | AB738264 | [45] |
| H287 | 0 | 0 | 1 | 0 | LC054400 | [51] |
| H288 | 0 | 0 | 1 | 0 | LC054401 | [51] |
| H289 | 0 | 0 | 1 | 0 | LC054402 | [51] |
| H290 | 0 | 0 | 1 | 0 | LC054403 | [51] |
| H291 | 0 | 0 | 1 | 0 | LC054404 | [51] |
| H292 | 0 | 0 | 1 | 0 | GBDCU003-12 | [52] |
| H293 | 0 | 0 | 0 | 1 | GBMNA38704-19 | [52] |
| H294 | 0 | 0 | 0 | 1 | MK403546 | [53] |
| H295 | 0 | 0 | 0 | 1 | GBMIN56951-17 | [46] |
| H296 | 0 | 0 | 1 | 0 | GBDP14515-13 | [52] |
| H297 | 0 | 0 | 1 | 0 | GBDP14517-13 | [52] |
| H298 | 0 | 1 | 0 | 0 | CULSA041-19 | http://www.boldsystems.org/ |
| H299 | 0 | 1 | 0 | 0 | CULSA043-19 | http://www.boldsystems.org/ |
| H300 | 0 | 0 | 1 | 0 | GBDP48513-19 | [45] |
| H301 | 0 | 0 | 1 | 0 | GBMIN56978-17 | [44] |
| H302 | 0 | 0 | 1 | 0 | GBMIN56979-17 | [44] |
| H303 | 0 | 0 | 0 | 1 | LRMQS002-17 | [47] |
| H304 | 0 | 0 | 0 | 1 | LRMQS035-17 | [47] |
| H305 | 0 | 0 | 0 | 1 | LRMQS066-17 | [47] |
| H306 | 0 | 0 | 0 | 1 | MLQSR050-17 | [47] |
| H307 | 0 | 0 | 0 | 1 | MLQSR051-17 | [47] |
| H308 | 0 | 0 | 0 | 1 | MLQSR055-17 | [47] |
| H309 | 0 | 0 | 0 | 1 | GBDP48491-19 | [48] |
| H310 | 0 | 0 | 0 | 1 | GBDP48493-19 | [48] |
| H311 | 0 | 0 | 0 | 1 | GBMIN56957-17 | [48] |
| H312 | 0 | 0 | 0 | 1 | GBMIN56958-17 | [48] |
| H313 | 0 | 0 | 0 | 1 | GBMIN56960-17 | [48] |
| H314 | 0 | 0 | 0 | 1 | GBMIN56965-17 | [48] |
| H315 | 0 | 0 | 0 | 1 | GBMIN56967-17 | [48] |
| H316 | 0 | 0 | 0 | 1 | GBMIN56969-17 | [48] |
| H317 | 0 | 0 | 0 | 1 | GBMIN56972-17 | [48] |
| H318 | 0 | 0 | 0 | 1 | GBMIN56974-17 | [48] |
| H319 | 1 | 0 | 0 | 0 | BBDIT1116-11 | http://www.boldsystems.org/ |
| H320 | 1 | 0 | 0 | 0 | BBDIT1121-11 | http://www.boldsystems.org/ |
| H321 | 1 | 0 | 0 | 0 | BBDIT1122-11 | http://www.boldsystems.org/ |
| H322 | 1 | 0 | 0 | 0 | BBDIT1123-11 | http://www.boldsystems.org/ |
| H323 | 1 | 0 | 0 | 0 | BBDIV1558-12 | http://www.boldsystems.org/ |
| H324 | 1 | 0 | 0 | 0 | GMGAA514-13 | http://www.boldsystems.org/ |
| H325 | 1 | 0 | 0 | 0 | GMGCC029-13 | http://www.boldsystems.org/ |
| H326 | 1 | 0 | 0 | 0 | GMGDD139-13 | http://www.boldsystems.org/ |
| H327 | 1 | 0 | 0 | 0 | IUP1014-14 | http://www.boldsystems.org/ |
| H328 | 1 | 0 | 0 | 0 | IUP1016-14 | http://www.boldsystems.org/ |
| H329 | 1 | 0 | 0 | 0 | IUP1018-14 | http://www.boldsystems.org/ |
| H330 | 1 | 0 | 0 | 0 | IUP675-14 | http://www.boldsystems.org/ |
| H331 | 1 | 0 | 0 | 0 | IUP885-14 | http://www.boldsystems.org/ |
| H332 | 1 | 0 | 0 | 0 | MOSN019-17 | http://www.boldsystems.org/ |
| H333 | 1 | 0 | 0 | 0 | MOSN044-17 | http://www.boldsystems.org/ |
| H334 | 1 | 0 | 0 | 0 | MOSN088-17 | http://www.boldsystems.org/ |
| H335 | 1 | 0 | 0 | 0 | MOSN1014-18 | http://www.boldsystems.org/ |
| H336 | 1 | 0 | 0 | 0 | MOSN1083-18 | http://www.boldsystems.org/ |
| H337 | 1 | 0 | 0 | 0 | MOSN1374-18 | http://www.boldsystems.org/ |
| H338 | 1 | 0 | 0 | 0 | MOSN1382-18 | http://www.boldsystems.org/ |
| H339 | 1 | 0 | 0 | 0 | MOSN1421-18 | http://www.boldsystems.org/ |
| H340 | 1 | 0 | 0 | 0 | MOSN1424-18 | http://www.boldsystems.org/ |
| H341 | 1 | 0 | 0 | 0 | MOSN1425-18 | http://www.boldsystems.org/ |
| H342 | 1 | 0 | 0 | 0 | MOSN1428-18 | http://www.boldsystems.org/ |
| H343 | 1 | 0 | 0 | 0 | MOSN1615-18 | http://www.boldsystems.org/ |
| H344 | 1 | 0 | 0 | 0 | MOSN1652-18 | http://www.boldsystems.org/ |
| H345 | 1 | 0 | 0 | 0 | MOSN179-18 | http://www.boldsystems.org/ |
| H346 | 1 | 0 | 0 | 0 | MOSN1869-18 | http://www.boldsystems.org/ |
| H347 | 1 | 0 | 0 | 0 | MOSN2115-19 | http://www.boldsystems.org/ |
| H348 | 1 | 0 | 0 | 0 | MOSN2116-19 | http://www.boldsystems.org/ |
| H349 | 1 | 0 | 0 | 0 | MOSN2480-19 | http://www.boldsystems.org/ |
| H350 | 1 | 0 | 0 | 0 | MOSN2487-19 | http://www.boldsystems.org/ |
| H351 | 1 | 0 | 0 | 0 | MOSN2647-19 | http://www.boldsystems.org/ |
| H352 | 1 | 0 | 0 | 0 | MOSN2649-19 | http://www.boldsystems.org/ |
| H353 | 1 | 0 | 0 | 0 | MOSN2655-19 | http://www.boldsystems.org/ |
| H354 | 1 | 0 | 0 | 0 | MOSN288-18 | http://www.boldsystems.org/ |
| H355 | 1 | 0 | 0 | 0 | MOSN2897-19 | http://www.boldsystems.org/ |
| H356 | 1 | 0 | 0 | 0 | MOSN2909-19 | http://www.boldsystems.org/ |
| H357 | 1 | 0 | 0 | 0 | MOSN2913-19 | http://www.boldsystems.org/ |
| H358 | 1 | 0 | 0 | 0 | MOSN292-18 | http://www.boldsystems.org/ |
| H359 | 1 | 0 | 0 | 0 | MOSN2953-19 | http://www.boldsystems.org/ |
| H360 | 1 | 0 | 0 | 0 | MOSN2978-19 | http://www.boldsystems.org/ |
| H361 | 1 | 0 | 0 | 0 | MOSN3117-19 | http://www.boldsystems.org/ |
| H362 | 1 | 0 | 0 | 0 | MOSN3126-19 | http://www.boldsystems.org/ |
| H363 | 1 | 0 | 0 | 0 | MOSN3127-19 | http://www.boldsystems.org/ |
| H364 | 1 | 0 | 0 | 0 | MOSN3136-19 | http://www.boldsystems.org/ |
| H365 | 1 | 0 | 0 | 0 | MOSN3165-19 | http://www.boldsystems.org/ |
| H366 | 1 | 0 | 0 | 0 | MOSN3182-19 | http://www.boldsystems.org/ |
| H367 | 1 | 0 | 0 | 0 | MOSN3228-19 | http://www.boldsystems.org/ |
| H368 | 1 | 0 | 0 | 0 | MOSN3318-19 | http://www.boldsystems.org/ |
| H369 | 1 | 0 | 0 | 0 | MOSN3319-19 | http://www.boldsystems.org/ |
| H370 | 1 | 0 | 0 | 0 | MOSN3321-19 | http://www.boldsystems.org/ |
| H371 | 1 | 0 | 0 | 0 | MOSN3381-19 | http://www.boldsystems.org/ |
| H372 | 1 | 0 | 0 | 0 | MOSN3402-19 | http://www.boldsystems.org/ |
| H373 | 1 | 0 | 0 | 0 | MOSN3403-19 | http://www.boldsystems.org/ |
| H374 | 1 | 0 | 0 | 0 | MOSN3490-19 | http://www.boldsystems.org/ |
| H375 | 1 | 0 | 0 | 0 | MOSN3499-19 | http://www.boldsystems.org/ |
| H376 | 1 | 0 | 0 | 0 | MOSN3523-19 | http://www.boldsystems.org/ |
| H377 | 1 | 0 | 0 | 0 | MOSN360-18 | http://www.boldsystems.org/ |
| H378 | 1 | 0 | 0 | 0 | MOSN436-18 | http://www.boldsystems.org/ |
| H379 | 1 | 0 | 0 | 0 | MOSN491-18 | http://www.boldsystems.org/ |
| H380 | 1 | 0 | 0 | 0 | MOSN503-18 | http://www.boldsystems.org/ |
| H381 | 1 | 0 | 0 | 0 | MOSN848-18 | http://www.boldsystems.org/ |
| H382 | 1 | 0 | 0 | 0 | MOSN852-18 | http://www.boldsystems.org/ |
| H383 | 1 | 0 | 0 | 0 | MOSN853-18 | http://www.boldsystems.org/ |
| H384 | 1 | 0 | 0 | 0 | MOSN858-18 | http://www.boldsystems.org/ |
| H385 | 1 | 0 | 0 | 0 | MOSN968-18 | http://www.boldsystems.org/ |
| H386 | 1 | 0 | 0 | 0 | MOSN972-18 | http://www.boldsystems.org/ |
| H387 | 1 | 0 | 0 | 0 | MOSN976-18 | http://www.boldsystems.org/ |
| H388 | 1 | 0 | 0 | 0 | MOSN991-18 | http://www.boldsystems.org/ |
| H389 | 1 | 0 | 0 | 0 | NEONT275-10 | [53] |
| H390 | 1 | 0 | 0 | 0 | NEONU128-11 | http://www.boldsystems.org/ |
| H391 | 1 | 0 | 0 | 0 | NEONU130-11 | http://www.boldsystems.org/ |
| H392 | 1 | 0 | 0 | 0 | NEONU309-11 | http://www.boldsystems.org/ |
| H393 | 1 | 0 | 0 | 0 | TDWGB127-10 | http://www.boldsystems.org/ |
| H394 | 1 | 0 | 0 | 0 | TDWGB705-10 | http://www.boldsystems.org/ |
| H395 | 1 | 0 | 0 | 0 | USDIQ1433-10 | http://www.boldsystems.org/ |
| Total | 1049 | 2 | 46 | 87 | 1184 |  |

**Figure 1.**
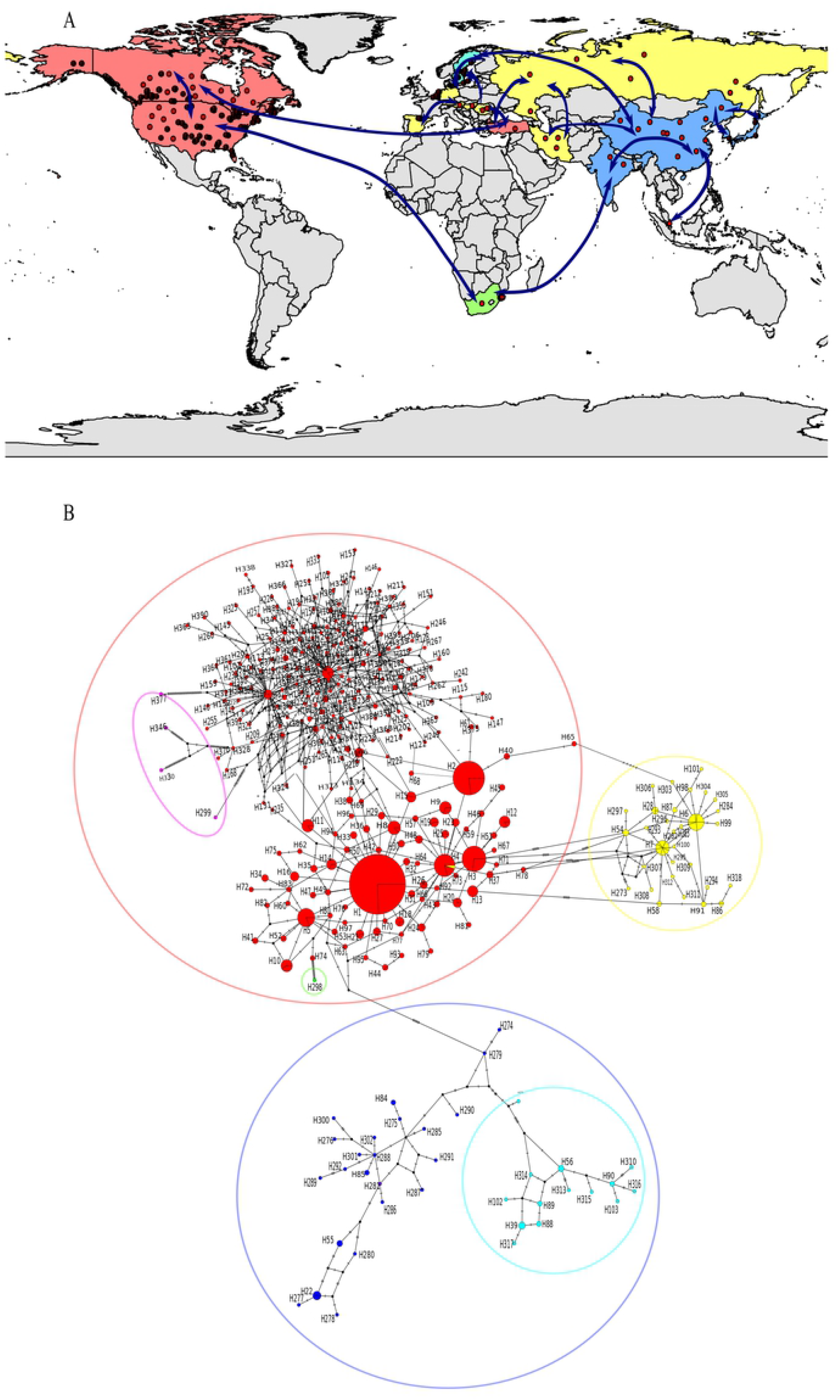
Locations where genetic material was extracted for the *A. vexans* populations analyzed (2A), as well as phylogenetic relations among populations, as arrows observed in the haplotype network (2B). In both figures the colors below denote the continents and their respective countries. 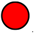 America + Europe: Canada, United States and Turkey. 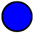 Asia: China, India, Japan, Singapore and South Korea. 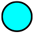 Europe + Asia: Sweden, Belgium and China. 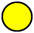 Europe + Asia: Romania, Sweden, Belgium, Russia, Kosovo, the Netherlands, China, Spain, Germany, Iran, Austria and Hungary. 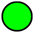 Africa: South Africa. 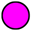 America + Africa: South Africa and the United States.

Table 2 shows, by continent and countries, the results of *Hd*, π and the different neutrality tests. In general, the global *Hd* was 0.92, while by continents it varied between 0.90 (Europe) and 1.0 (Africa). The *Hd* by countries was between 0.0 (Austria, Turkey, Hungary, Singapore, and India) and 1.0 (Germany, Kosovo, Rumania, Russia, South Korea, and South Africa). In turn, the global π was 0.01, while by continents it varied between 0.005 (Europe) and 0.08 (Africa). The π among countries ranged between 0.0 (Austria, Hungary, Turkey, Singapore, and India) and 0.08 (South Africa). Neutrality tests, Tajima’s D, and Fu’s F, at global level, were negatively significant (D = −2.20, p < 0.001; F = −5.22, p < 0.02); by continent and countries, America (D = −2.46, p < 0.05; F= −5.65, p < 0.02) and all its countries (Canada D = −2.43, p < 0.05; F = −5.13, p < 0.02 and USA D = −2.30, p < 0.05; F = −4.96, p < 0.02) were statistically significant and with negative values.

**Table 2.**
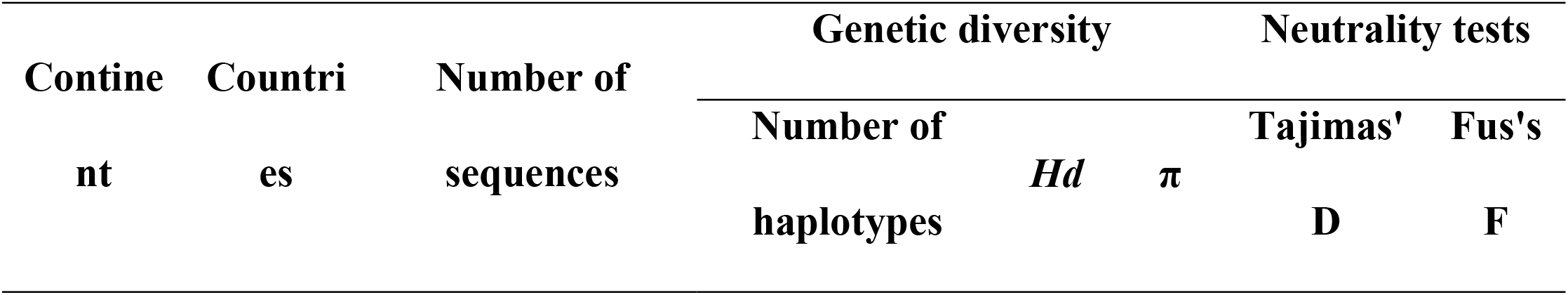
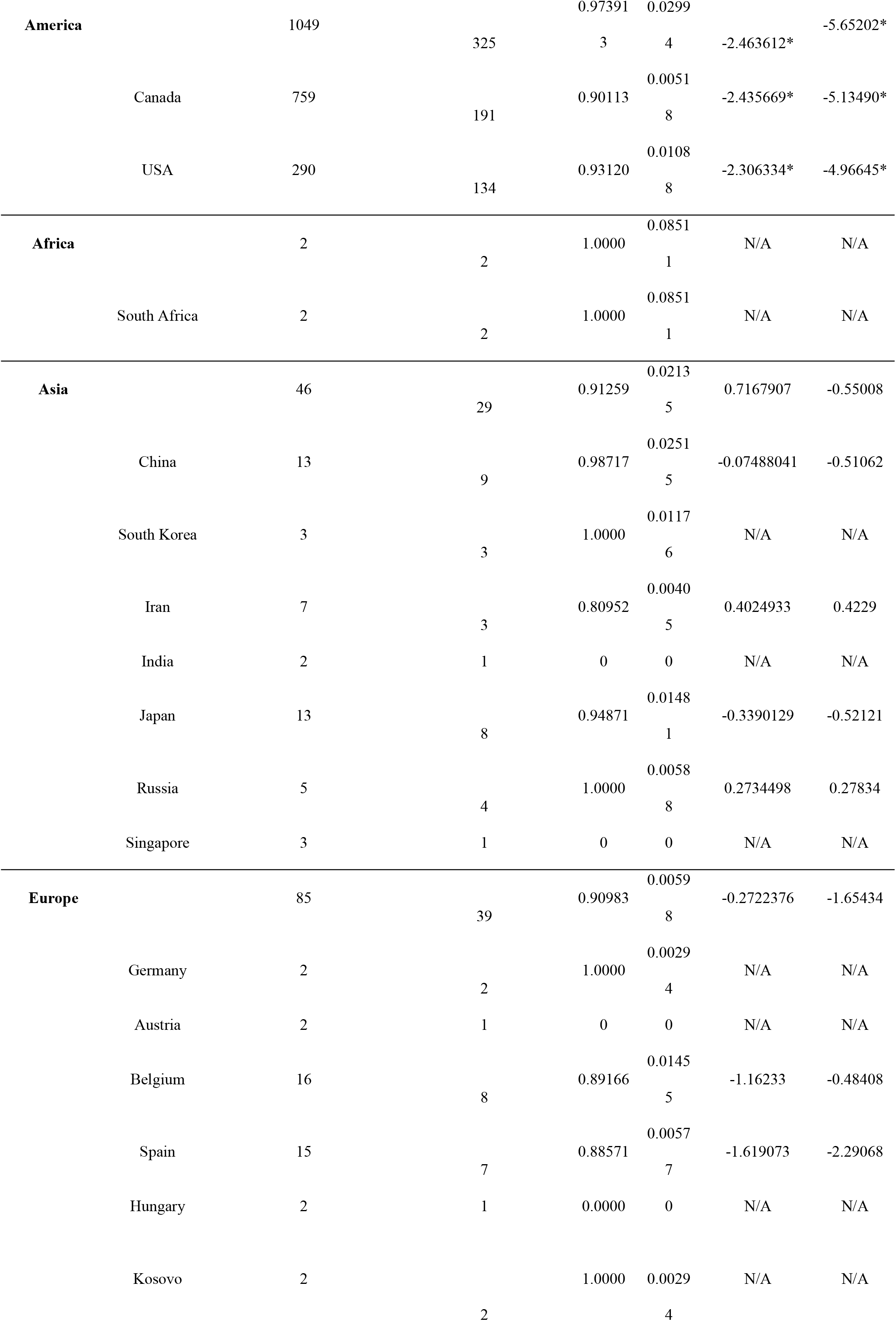
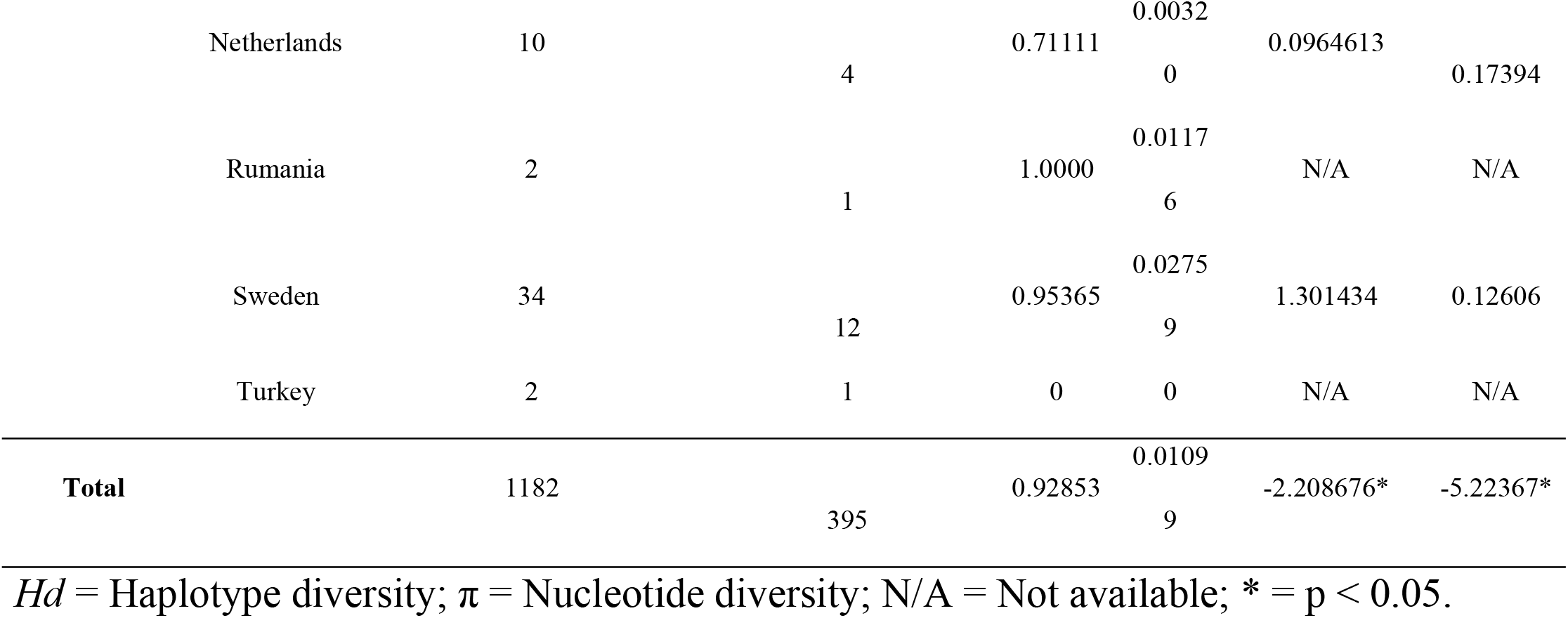
Results of genetic diversity and neutrality tests at global level and by countries for *A. vexans*.

In Table 3, the AMOVA indicated the existence of genetic structuring at continent and country levels and within countries (F_ST_ = 0.08, p < 0.05), where the highest variation percentage was observed among *A. vexans* individuals within countries (91.30%), followed by 7.07% among continents and 1.62% among countries in the same continent. In Figure 2, Mantel’s test indicated no isolation by distance (r = 0.003, p > 0.05).

**Table 3.**
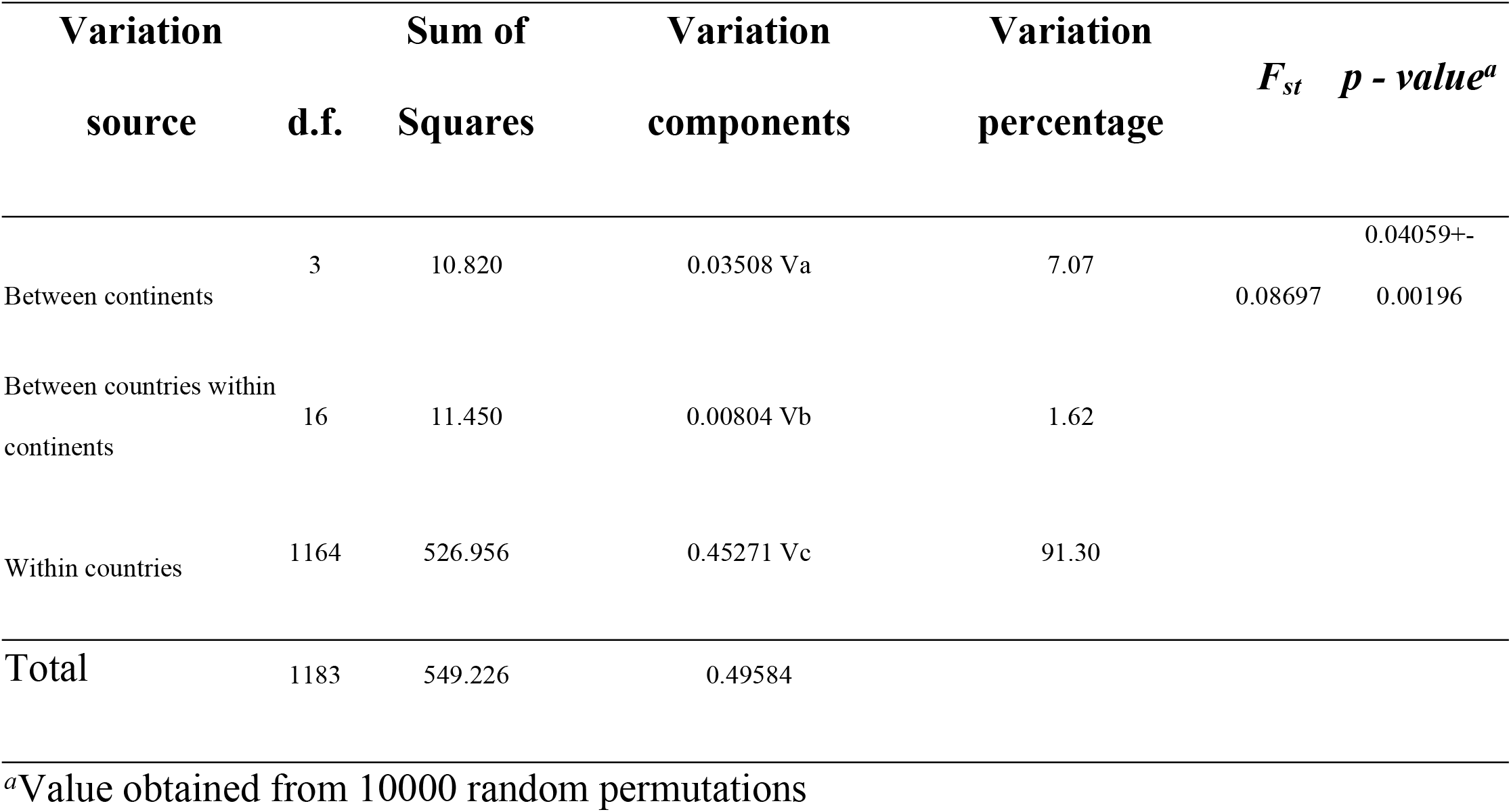
Analysis of molecular variance (AMOVA) of populations of *A. vexans* at continental level, by countries, and within them.

**Figure 2.**
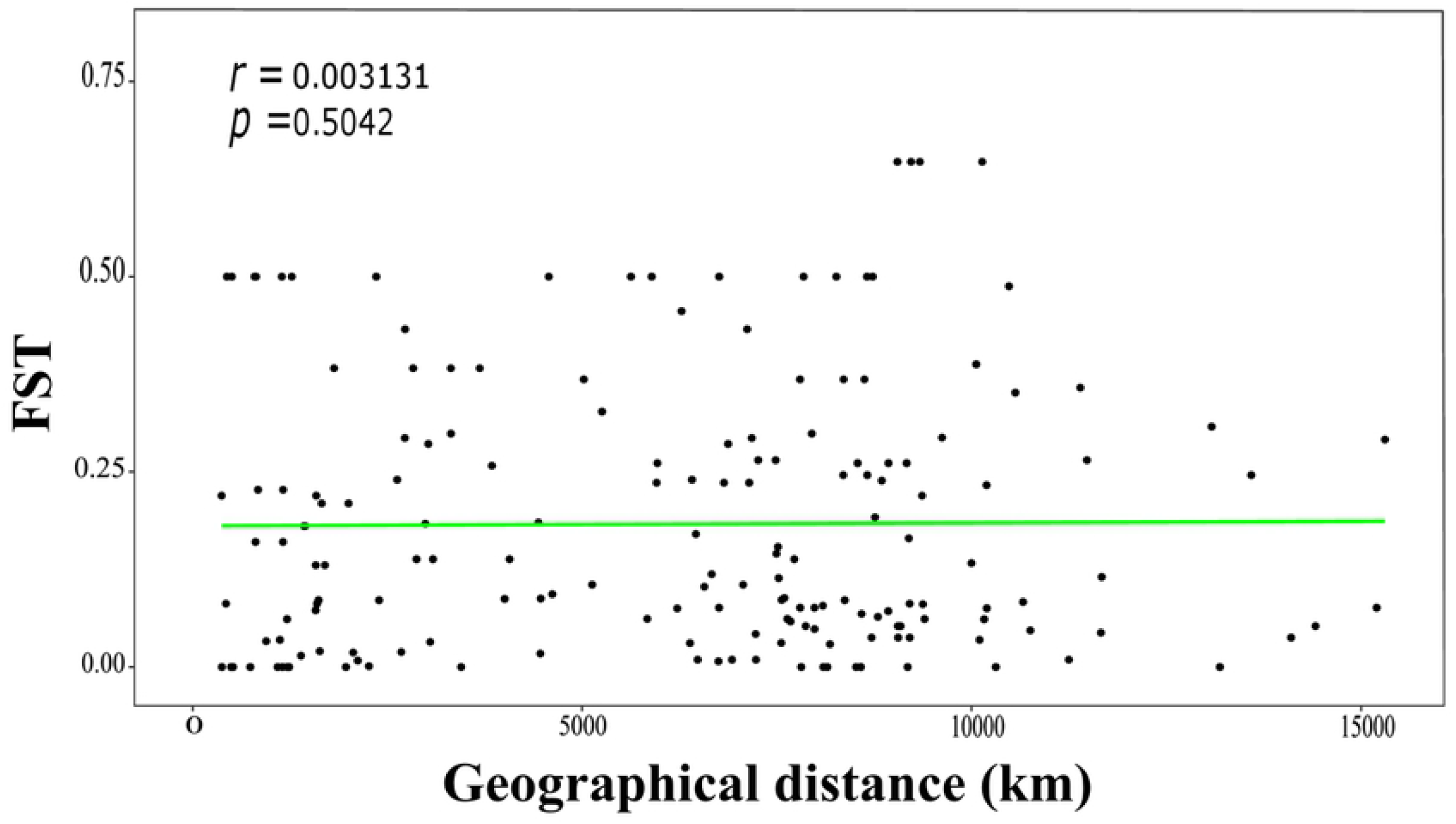
Global Mantel’s correlation test for genetic and geographic distance of the *A. vexans* populations analyzed.

Table 4 shows peer-to-peer comparisons among countries after Bonferroni’s correction. For these, significant genetic structuring was detected within Canada with respect to Belgium, Sweden, Spain, Hungary, and Netherlands. The USA with respect to Austria, Belgium, Sweden, Spain, Hungary, and Netherlands. China with respect to Netherlands, the USA, and Canada. India with respect to the USA and Canada. Iran with respect to the USA and Canada. Japan with respect to Belgium, Sweden, Spain, and Netherlands. Russia with respect to the USA and Canada. Singapore with respect to Belgium, Sweden, and Spain.

**Table 4.**
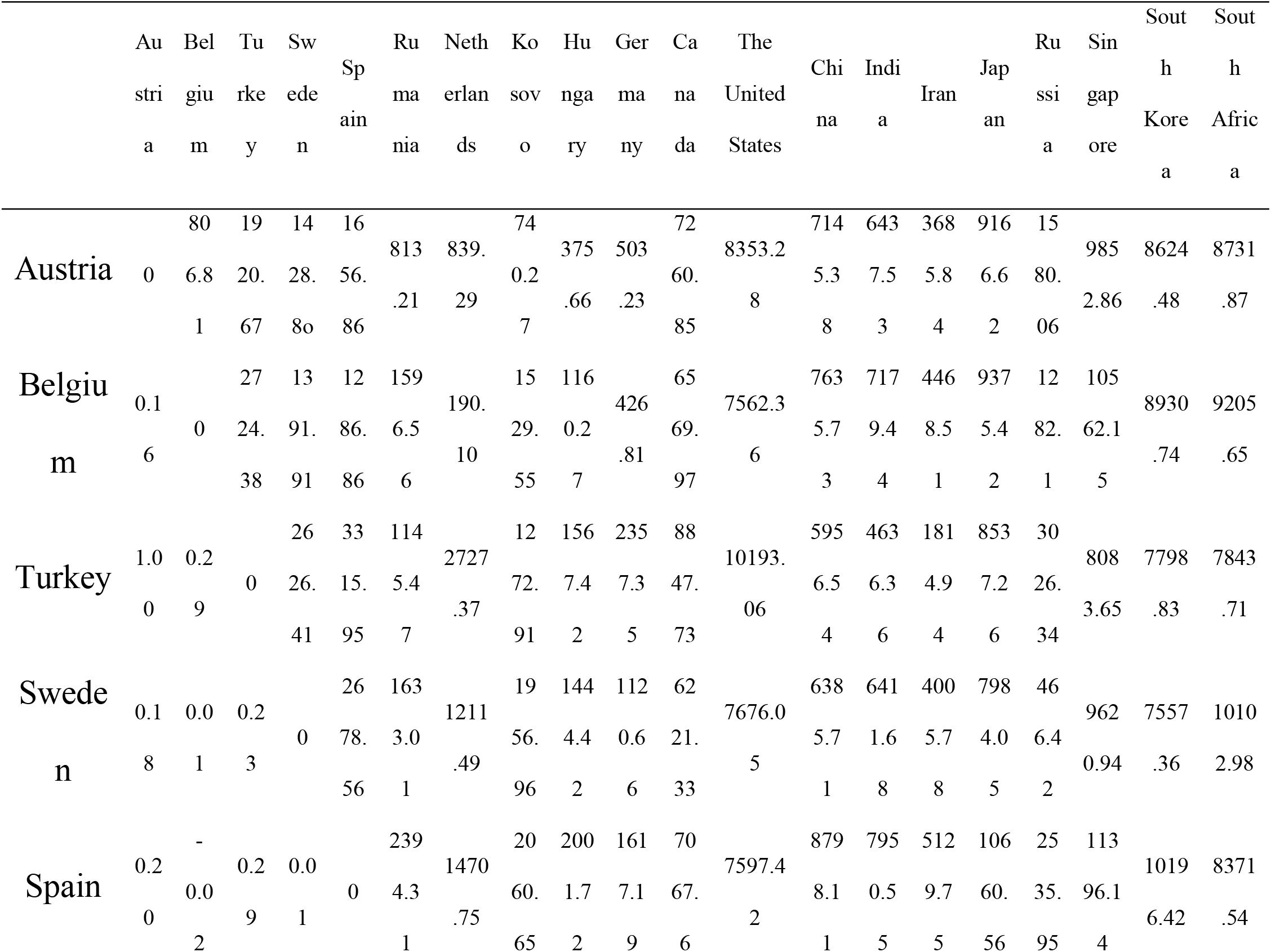
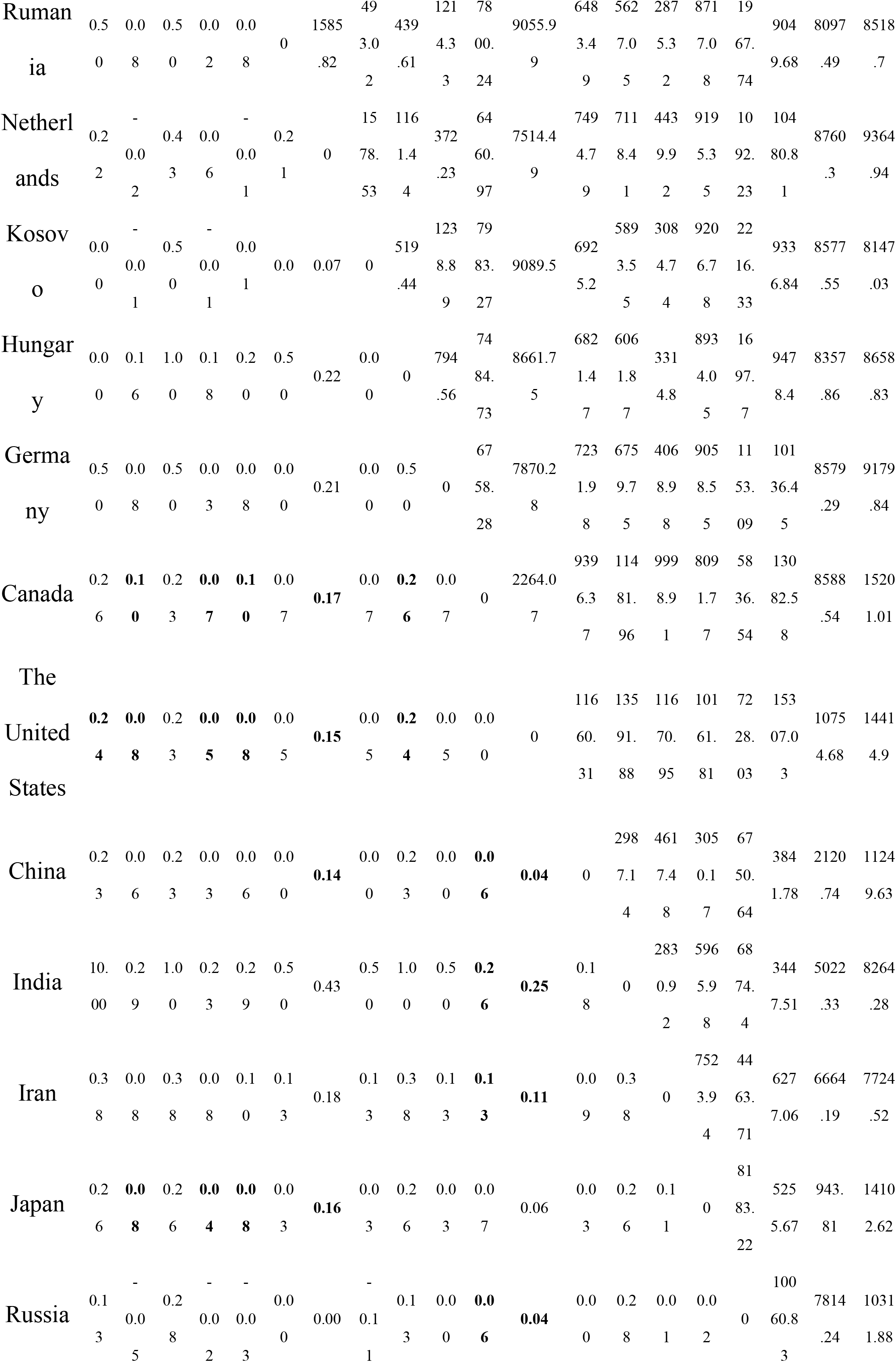
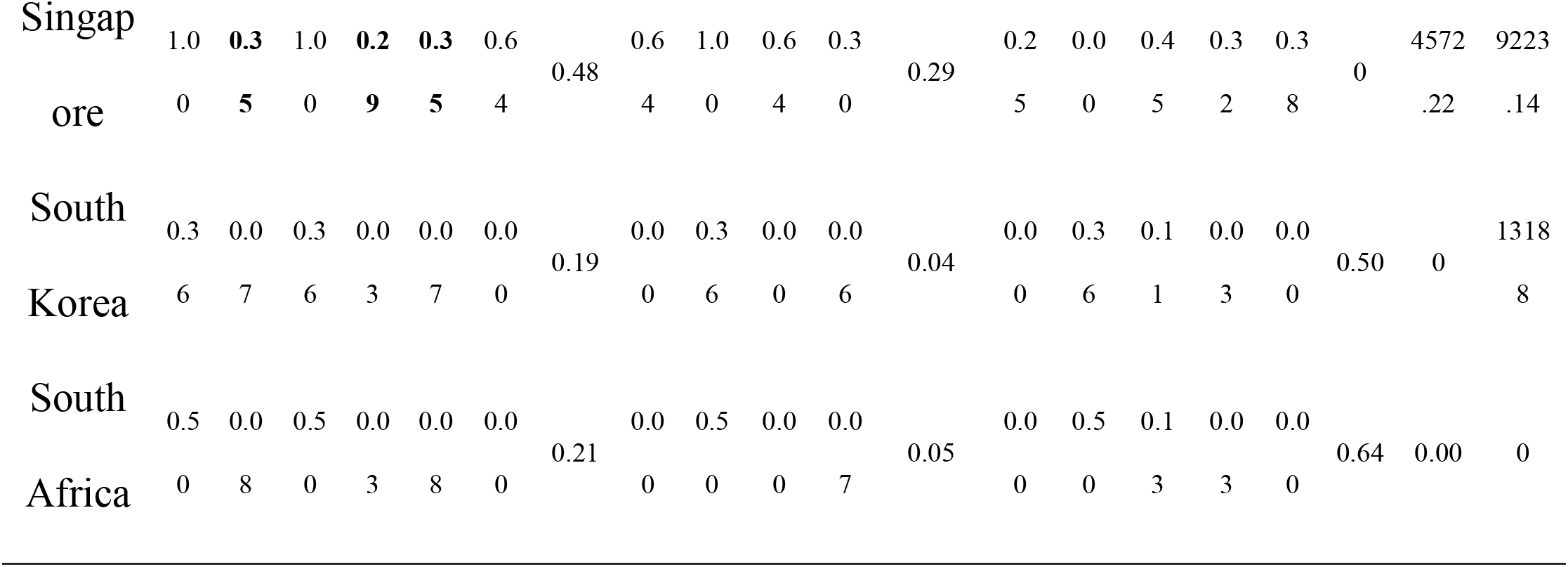
Values by countries of genetic differentiation by peers (F_ST_) and geographic distance (Km) among *A. vexans* populations. Values in bold are genetically structured populations.

Figure 3 shows the results of the phylogenetic analyses by using BI (Fig. 3A) and ML (Fig. 3B); both analyses recovered the same six clades, but with distinct topologies: clade I grouped mosquito populations from the United States, Canada, and Turkey (n = 322 H); clade II, populations from China, Japan, Singapore, and South Korea (n = 23); clade III, populations from Sweden, Belgium, and China (n = 14 H); clade IV, mosquito populations from Rumania, Sweden, Belgium, Russia, Kosovo, Netherlands, China, Spain, Germany, Iran, Austria, and Hungary (n = 31 H); clade V, populations from South Africa (n = 1 H); and clade VI, populations from the United States and South Africa (n = 4 H). Similar results were observed in the haplotype network (Figure 1B).

**Figure 3.**
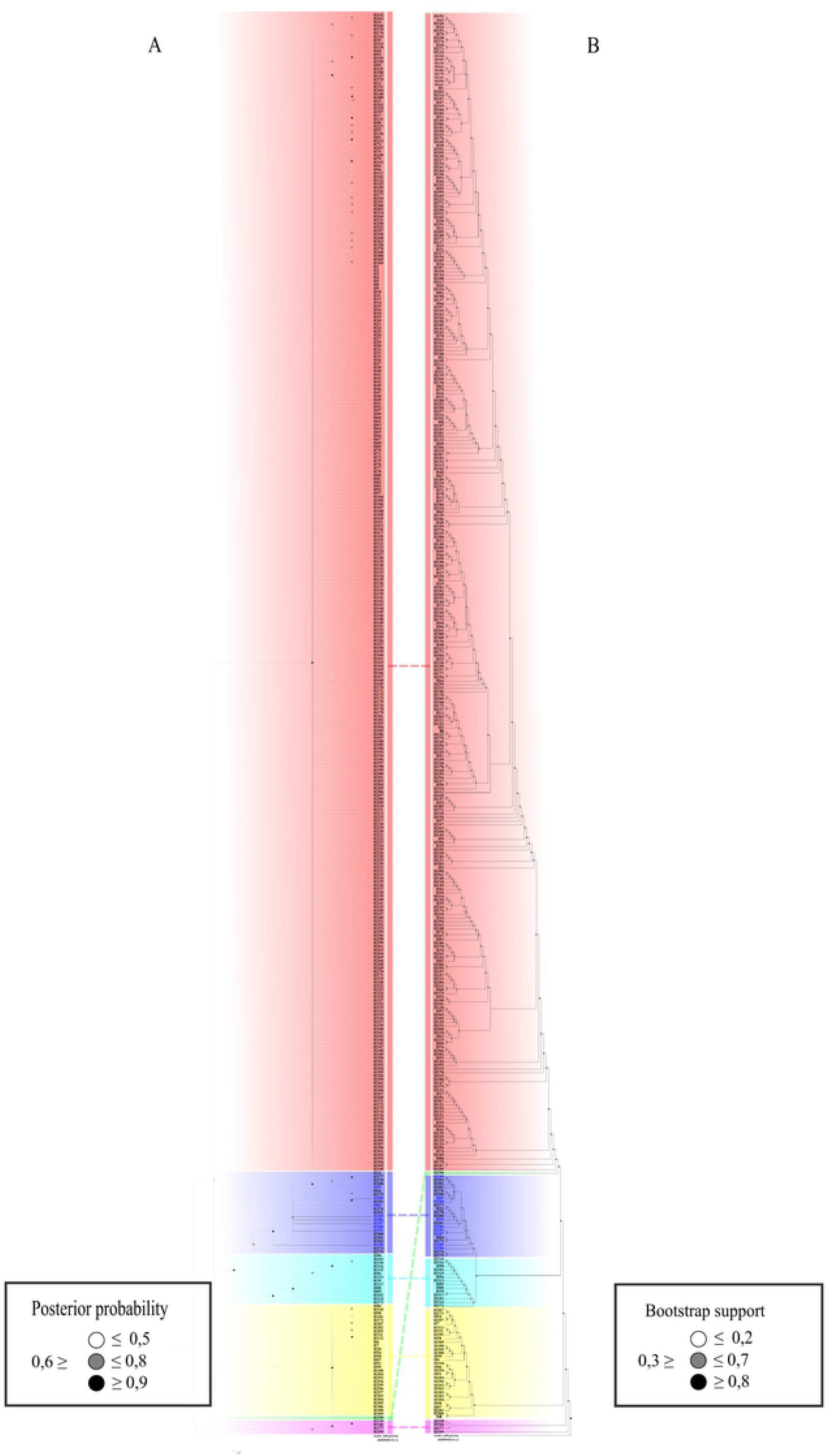
Phylogenetic tree for *Aedes vexans* populations constructed from 395 haplotypes from the COI gene by using Bayesian Inference, BI, (3A) and Maximum Likelihood, ML, (3B). The evolutionary history for both analyses was inferred by using the GTR + G model, as suggested by jModelTest version 2.1.10. The BI tree was obtained by using 2-million generations, while the ML used 1,000 replicas. For BI, the support of the branches is indicated by the subsequent probability values, while for ML the bootstrap values are shown. Numbers in blue represent sequences from the *A. nipponii* subspecies. In both figures the colors below denote the continents and their respective countries. 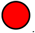 America + Europe: Canada, United States and Turkey. 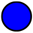 Asia: China, India, Japan, Singapore and South Korea. 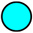 Europe + Asia: Sweden, Belgium and China. 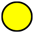 Europe + Asia: Romania, Sweden, Belgium, Russia, Kosovo, the Netherlands, China, Spain, Germany, Iran, Austria and Hungary. 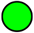 Africa: South Africa. 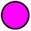 America + Africa: South Africa and the United States.

## Discussion

To our knowledge, this is the first study on the genetic structure of *A. vexans* using all the genetic information available for the COI gene from the GenBank and Boldsystem databases. From this information, both the haplotype network and the phylogenetic tree revealed the existence of six clades; clade I grouped mosquito populations from America and Europe; clade II, populations from Asia; clades III and IV, populations from Europe and Asia; clade V, populations from Africa; and clade VI, populations from America and Africa. For *A. vexans*, existence is suggested of three subspecies: *A. vexans vexans, A. vexans arabiensis*, and *A. vexans nipponii*, from which it was possible to include in our natural population analyses of *A. vexans* and *A. vexans nipponii. A. vexans vexans* has been reported for east Asia and Oceania, *A. vexans arabiensis* in Africa and Europe, and *A. vexans nipponii* in southeast Asia [1,14,2,42,15,16,17]. However, our results do not suggest the existence of three subspecies or the geographic relations observed. Nonetheless, it is interesting that with *A. vexans* being considered native of America [13] its invasion is not hypothesized to other latitudes, a pattern observed even with *A. vexans nipponii* terminals registered in Asia, as suggested by our results. This is why the subspecies observed in countries different from America would probably be populations from their place of origin through passive transport (*i*.*e*., maritime, air, or land transport), as already observed for other Culicidae invaders [54,55,56,57,58], including *A. vexans* [59]. Future studies should include information at genome level to try to solve the possible existence of subspecies, given that using a single marker is not sufficient to define the species [60], as was observed even for this species by using the COI gene [48].

Haplotype diversity and number of haplotypes observed in the American continent (*Hd* = 0.97; H = 325) were higher than in other continents; for example, Europe (*Hd* = 0.90; H = 39). Various studies suggest that native species have higher genetic diversity when compared with places different from their native area [61,62,63]. For example, a global study on the Cosmopolitan Asian mosquito, *Aedes albopictus*, principal dengue, Zika and Chikungunya vector in Asia and Europe, observed greater genetic diversity (*Hd* = 0.94, π = 1.60) in its native area with respect to all the areas it has invaded, showing – therein – lower diversity indices (lower *Hd* in Netherlands (0.059 = π = 0.011) and higher *Hd* in China (0.946 = π = 1.609)) [63]. The aforementioned supports our hypothesis that *A. vexans* populations may have invaded other latitudes.

The *A. vexans* mosquito showed significant genetic divergence among some populations from the American continent with respect to some European and Asian populations (Table 4). [64], analyzing natural *A. vexans* populations from the United States and Germany, found that these do not share a common gene pool, proposing that the geographic barriers formed by the Atlantic and Pacific Oceans impede gene flow and cause genetic changes in the evolutionary lineages of *A. vexans*. However, our results suggest no existence of geographic and genetic isolation. Additionally, for most of the populations, the results of the neutrality tests, Tajima’s D and Fu’s FS were negative (Table 2), suggesting that these have experienced recent bottlenecks and population expansion [65]. This may be due to recent vector-control actions and colonization events, phenomena commonly observed in mosquitos of medical and veterinary importance [66]. In the first case, these are used to diminish the population size of *A. vexans* and other vector species of diseases and, consequently, curtail the epidemiologic transmission of the diseases it transmits [67,68,69]. In the second case, recent colonization events may take place in areas from where the *A. vexans* populations are lost during harsh winters or after vector-control actions and can be re-colonized by surviving individuals from neighboring areas [3,70,58].

## Conclusions

Finally, our results suggest that the *A. vexans* populations that invaded other continents originate directly from America, where possibly transcontinental commercial routes favored their long-distance dispersion. Moreover, we consider this study as the base for future taxonomic research that address the problem of the existence of subspecies within *A. vexans*, given that our results did not recover any of the subspecies suggested by the literature.

## Acknowledgments

The authors thank the Vice-rectory of Research at Universidad del Quindío for the funding needed to translate the manuscript.

## Ethical considerations

This work did not experiment with humans or other living beings; its data were obtained from genetic databases freely available on line.

## Conflicts of interest

The authors have no conflict of interest to declare.

## Supporting information

**S1**. Nucleotide diversity of the COI gene for A. vexans populations. Mafft alignment to compare the haplotypes from this study. The amino acid sequence translated is represented by capital letters in blue, over the first nucleotide of its corresponding codon. Invariable sites are indicated with points, contrary to the alternative nucleotide, and spaces with (-).

